# Impact of Variability in Cell Cycle Periodicity on Cell Population Dynamics

**DOI:** 10.1101/2021.10.13.464184

**Authors:** Chance M. Nowak, Tyler Quarton, Leonidas Bleris

**Author notes:** Correspondence should be addressed to L.B.

## Abstract

The cell cycle consists of a series of orchestrated events controlled by molecular sensing and feedback networks that ultimately drive the duplication of total DNA and the subsequent division of a single parent cell into two daughter cells. The ability to block the cell cycle and synchronize cells within the same phase has helped understand factors that control cell cycle and the properties of each individual phase. Intriguingly, when cells are released from a synchronized state, they do not maintain synchronized cell division and rapidly become asynchronous. The rate and factors that control cellular desynchronization remain largely unknown. In this study, using a combination of experiments and simulations, we investigate the desynchronization properties in cervical cancer cells (HeLa) starting from the G_1_/S boundary following double-thymidine block. Propidium iodide (PI) DNA staining was used to perform flow cytometry cell cycle analysis at regular intervals of 8 hours, and a custom auto-similarity function to assess the desynchronization and quantify the convergence to asynchronous state. In parallel, we developed a single-cell phenomenological model the returns the DNA concentration across the cell cycle stages and fitted the parameters using experimental data. Simulations of population of cells reveal that the cell cycle desynchronization rate is primarily sensitive to the variability of cell cycle duration within a population. To validate the model prediction, we introduced lipopolysaccharide (LPS) to increase cell cycle noise. Indeed, we observed an increase in cell cycle variability under LPS stimulation in HeLa cells, accompanied with an enhanced rate of cell cycle desynchronization. Our results show that the desynchronization rate of artificially synchronized in-phase cell populations can be used a proxy of the degree of variance in cell cycle periodicity, an underexplored axis in cell cycle research.

## Introduction

Cell division is traditionally described as a general process divided into two phases, the interphase and mitosis (cell division). Interphase is further divided into three subphases; Gap 1 phase (G_1_) in which the cell has a DNA content of 2N, synthesis phase (S) in which the cell’s DNA content is greater than 2N but less than 4N, and Gap 2 phase (G_2_) in which the cell’s DNA content is 4N upon completion of synthesis. Early observations into cell cycle progression showed that the timing of G_1_ phase is highly variable not just between cell types but also between cells within a monoclonal population, and that this variable length directly impacts the heterogeneity observed in clonal populations for cell cycle periodicity ^1,2^. Additionally, a critical point in the cell cycle was discovered^3^, in which cells were found to be committed to DNA synthesis independent of environmental factors. Moreover, it was later demonstrated that under various suboptimal nutritional conditions, cell cycle progression could be arrested at the G_1_/S boundary, and escapement into S-phase could only occur once suitable nutritional needs were restored^4^. The boundary was termed the restriction point (R-point), whereby cells could enter a lower metabolic rate (a quiescent state) to remain viable until adequate nutrition is restored allowing the necessary constituents to be present in suitable amount to enable DNA synthesis^4^. Ultimately, it was shown that the high variability of G_1_ phase duration can be attributed to a cell’s ability to overcome the restriction point^5^.

Investigations into cell cycle progression and regulation often start with the need to synchronize cells within a population to the same cell cycle phase^6,7^. One common approach to cell cycle synchronization is the double-thymidine block that interferes with nucleotide metabolism resulting in an inability of the cells to synthesize DNA causing a cell cycle arrest at the G_1_/S boundary ^8,9^. Interestingly, when synchronized cell populations are released from cell cycle arrest, they quickly desynchronize, and reach a state of “asynchronicity,” whereby the individual cell cycle phases stabilize into fixed percentages within the overall population. Indeed, simply sampling cells from an asynchronously growing *in vitro* cell culture will reveal (**Figure 1a**) the fixed percentages for the three phases of interphase (G_1_, S, and G_2_). Additionally, cells can be pulse-labeled with bromodeoxyuridine (BrdU) to create a semi-synchronous cell population in which only cells in actively progressing through S-phase incorporate the thymidine analog BrdU into their genome, and thus the original pulse-labeled population can be tracked overtime by using a fluorescently conjugated BrdU antibody^10^. These observations again showed that the initially pulse-labeled cells progressed synchronously through the cell cycle for some time before quickly desynchronizing and resorting back to an asynchronous DNA distribution profile.

**Figure 1:**
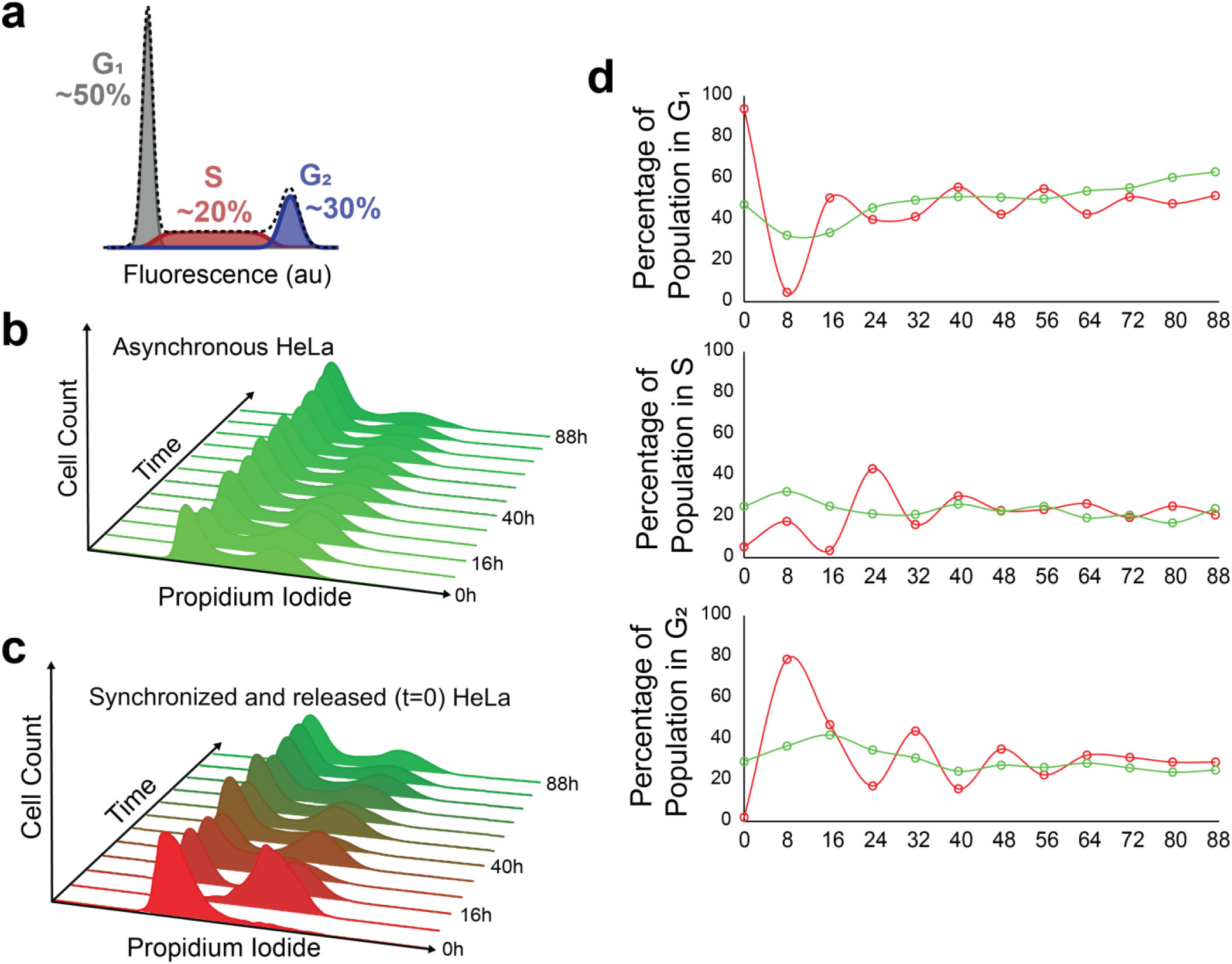
Cell Desynchronization via Double Thymidine Block and Release. **a)** Cell cycle phases as indicated by cell DNA content and approximate phase distribution in an asynchronous population. **b)** Fluorescent profile of propidium iodide (PI) stained cells during asynchronous growth from t=0 to t=88. **c)** Fluorescent profile of PI-stained cells following G_1_/S synchronization by double thymidine block from t=0 to t=88. **d)** Percentages of cells in a given cell cycle phase at a given time point; asynchronous cell growth in green and desynchronous cell growth in red. The cell cycle phase percentages for each time point were determined via the Dean-Jett-Fox model.

The inherent variability of cell cycle duration between identical cells may be accounted for by considering sources of cellular noise. In other words, the variability between cellular constituents such as signaling and transcriptional factors, along with the biochemical stochasticity of molecular interactions do likely propagate to the phenotypic level and may be responsible for varying timing events that dictate cell cycle progression. For example, signalling factors in a tumor microenvironment that confer a higher degree of intercell variability contribute to tumor cell heterogeneity and pathology^11,12^. Therefore, it is important to examine the implications of cellular noise to cell cycle periodicity.

In this report, we investigated the rate of cell cycle desynchronization by measuring the change in the DNA distribution of a population of cells over time. To this end, we measured the single-cell DNA concentration of a population of cells as they transition from an initial state of cell cycle synchrony, where cells are experimentally locked into the G_1_/S boundary, to a state of asynchrony. We used statistical tools to quantify the dynamic change in the DNA probability density function over time from an initial synchronized cell population. Subsequently, we developed a mathematical model to simulate at single-cell level the DNA concentration as the cell transitions through cell cycle states, and finally, experimentally validated our model prediction. More specifically, our model revealed that cell cycle desynchronization rates were particularly sensitive to the variability of cell cycle duration within a population. With this insight, to validate the results we introduced external noise in synchronized cells using lipopolysaccharide and, indeed, confirmed an increase in cell cycle desynchronization.

Considering the ubiquitous role of the cell cycle properties to cell health, the implications of our work extend to numerous fronts.

## Results

### Thymidine-based arrest and desynchronization

The exogenous introduction of excessive thymidine into cells interrupts DNA synthesis, arresting the population of cells in the G_1_/S-phase transition. Upon release, the population of cells are permitted to reenter their respective cell cycles. Ultimately, the population of cells will become asynchronous with respect to their cell cycles, yielding a PI fluorescent profile. The PI distributions dynamically change as the population desynchronizes.

After cells were synchronized via double-thymidine block, timepoints were collected every 8 hours for a total of 88 hours. Both asynchronous (untreated) cells (**Figure 1b**) and synchronized (**Figure 1c**) were subjected propidium iodide staining and flow cytometry analysis. Notably, we observed near full synchronization of cells as judge by the first few timepoints (**Figure 1c**) in the synchronous population. While inhibition of DNA synthesis can cause replicative errors due to stalled replication forks, resulting in quiescence or cell death, we did not observe neither an increase in cell death nor any quiescent populations, which would manifest as a sub-G_1_/G_1_ population at timepoint 8. Each PI histogram was subjected to cell cycle phase classifier^13–15^ with the cell cycle phase distribution displayed as percentages of the total population. As we observe in **Figure 1d**, the synchronized population eventually reaches an asynchronous distribution. The residual plots of the DNA distribution of the synchronous population against the asynchronous population ultimately converges to within 8.4%, 1.5%, and 6.1% of G_1_, S, and G_2_, respectively (**Supplementary Figure 1**).

### Quantifying cell synchronicity

The DNA dynamics during interphase of a population of cells is defined by the population’s collective distribution of its DNA at a given time. If all the cells within a population are undergoing interphase synchronously, time separated measurements of the population’s DNA distribution will accordingly change in time. This would mean that the DNA distribution of a population of cells will be different for each time measurement. Conversely, if the population’s cells are independently progressing through interphase, temporal differences between the population’s DNA distribution become indistinguishable, rendering its DNA distribution into a static steady-state (**Figure 1a**).

With this in mind, we can create a set of assumptions: Let {**X**_*t*_} denote sets of observations generated from an evolving probability distribution at any point in time *t*. We define the auto-similarity function (ASF) between times *t*_1_ and *t*_2_ as

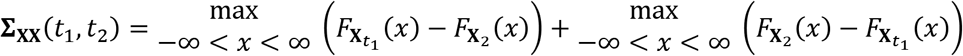

where *F*_**X**_*t*__ denotes the cumulative distribution function of a given set of observations **X**_*t*_.

Essentially, the auto-similarity function is the Kuiper two-sample test statistic, which measures the similarity between two sets of data, performed on a single, time evolving variable **X**_*t*_ rather than two distinct variables. The Kuiper test statistic is rotation-invariant, making its application insensitive to the “starting points” of the data to be compared. As the DNA content measured in our cell populations cycle between 2N to 4N, the data collected from our cell cycle experiments are inherently cyclical, making the use of a rotation-invariance test statistic ideal. If the evolving distribution eventually converges to a steady-state, we expect **∑_XX_**(*t*_*i*_, *t*_*i*+1_) → 0 for some successive time measurements *t_i_* and *t*_*i*+1_ as *t* → ∞, where a value of 0 indicates full asynchrony. Conversely, we interpret non-zero, positive evaluations of the ASF to indicate dissimilarity, where, in the case of a cyclically evolving sets of data, evidence that the underlying probability distribution is in a transient state, where a maximum value of 1 indicates full synchrony (**Figures 2a**).

**Figure 2:**
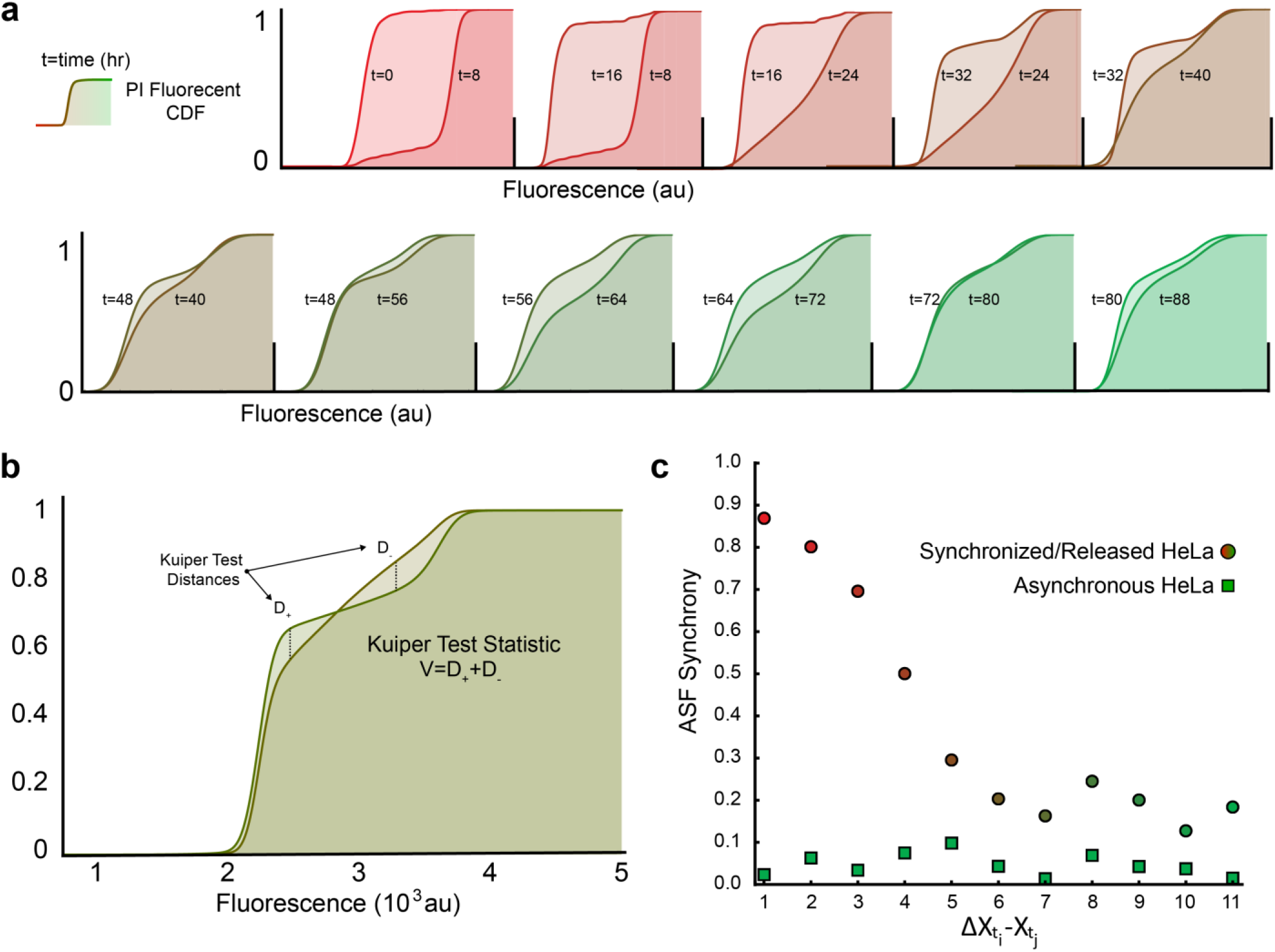
Rate of desynchronization using Kuiper Test Statistic. **a)** Pairwise comparison of PI CDFs for each time point (data shown is from synchronized cells). **b)** Visual representation of Kuiper Test Statistic determination between time points. **c)** Rate of desynchronization between asynchronous (green) and synchronized (red) Hela cells. Over time (~60 hours) synchronized cells being to reach an asynchronous state.

In our experiments, {**X**_*t*_} is variable DNA fluorescently measured by flow cytometry in PI-stained populations of cells, where *t*_*i*_ = {0,8,16,…,88} indicates the hour corresponding to the *i_th_* measurement of data collected with respect to their release from cycle arrest via double thymidine block at *t*_0_ = 0. We expect that the ASF evaluation of times *t*_0_ and *t*_1_ to be the greatest as the population of cells synchronously progress through the cell cycle, resulting in markedly dissimilar distributions of DNA in observation sets **X**_t_0__, and **X**_t_1__.

As the individual cells within a population variably progress through the cell cycle, we expect population DNA distributions to diverge, eventually settling to the classic asynchronous distribution profile (**Figure 1a-d**), where successive measurements of a no-longer-evolving variable are expected to be near-zero. We calculated the ASF between each temporally successive pair of data for both the synchronized cell population and the asynchronous control population (**Figure 2c**). We found that the ASF converges to a minimum of 0.127 from an initial value of 0.869, following a logistics curve. We observed an expected linear ASF from the asynchronous population with slight oscillations, most likely emerging from unintended loss of mitotic cells during harvesting (mitotic shake off) positive slope (**Figure 2c**).

### A Single Cell Interphase Model

Cell cycle progression is intimately linked to a cell’s dynamically changing DNA content. Temporal transitions from a cell’s state of 2N to 4N define cell cycle phases, where G_1_, S, and G_2_, correspond to genetic quantities of 2N, 2N+, and 4N, respectively, where the event of mitosis restarts the cell cycle for two progeny cells. Deterministically, we model a single cell’s dynamic DNA content as

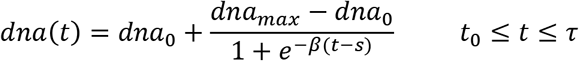

where *dna*_0_ is the initial genetic content in phase G_1_, *dna_max_* is the maximum genetic content after synthesis, *β* parameterizes the synthesis rate, *s* is the time in which the cell is halfway through synthesis and determines the periods of G_1_, S, and G_2_, and *t* is time. We assume that synthesis faithfully duplicates the genetic content, where *dna_max_* = 2 *dna_0_*, thus reducing the above equation to:

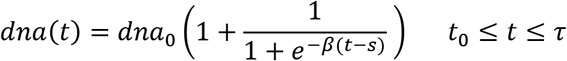

We can further reduce Eq. 2 by representing *β* and *s* as functions of the cycle period *τ* as:

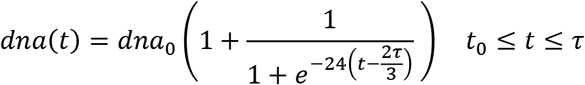

where we assume that the duration of S is ~1/3 of the total cycle period. Accordingly, this single cell model captures DNA concentration during interphase using two parameters, the initial DNA concentration and the cell cycle period (**Figure 3a**).

To study the impact of the cell cycle period to the rate of asynchrony we use the Error-in-Variables (EIV) modeling approach to add noise to the cycle periodicity:

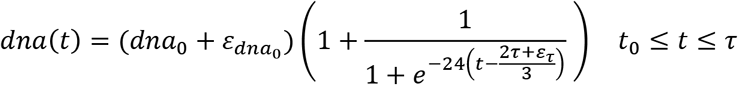

where 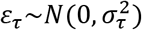 is a normally distributed error term with variance 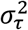 and 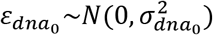 is additionally added to capture fluorescent variability seen as broadened peaks around G_1_ and G_2_ (**Figure 1a**). We simulate a population of 1,000 cells, each starting synchronously at G_1_ with extrinsically varying initial DNA content and cell cycle periodicity, as they repetitively progress through interphase (**Figure 3b**). We then take temporal slices of the DNA content of the population of cells and plot the populations distribution of DNA content intermittently (**Figure 3b-c**). We finally apply ASF to the slices in a pairwise manner as done in the experiment (**Figure 2**). An observation from these simulations is that a Poisson distributed error term for variance, as opposed to normal distribution, did not accurately capture the DNA dynamics of desynchronized cells using our ASF analysis.

**Figure 3:**
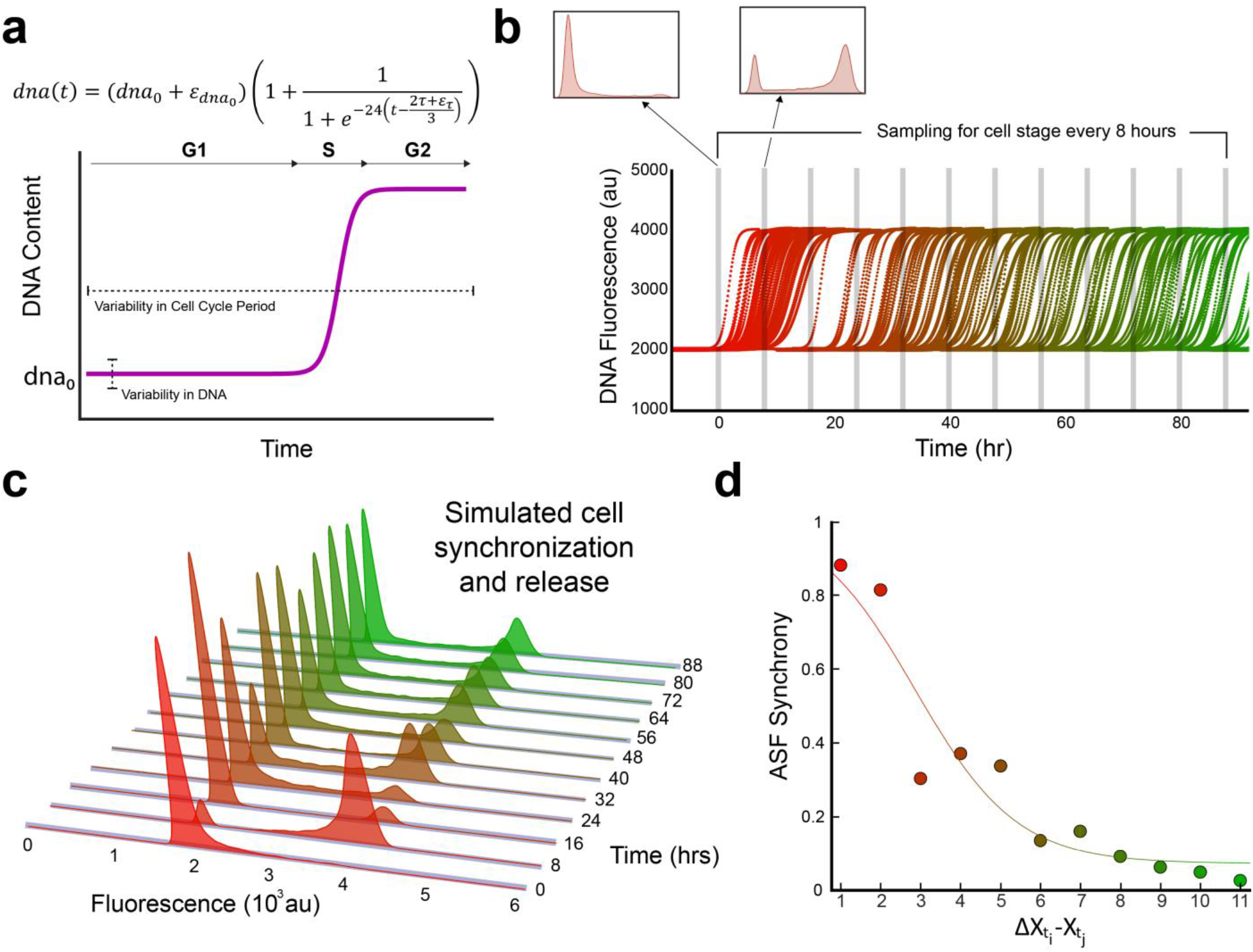
Single cell model of desynchronization. **a)** DNA synthesis is captured by the Gaussian error function where the relative durations of cycle phase are tunable. **b)** Cell cycle pace inheritance following a Gaussian distribution. **c)** Simulated data of PI staining of multiple lineages with normally distributed initial gene content. **d)** Desynchronization rate of simulated cell population.

Importantly, we found that only by including a variance term to cell cycle periodicity were we able to capture population dynamics that recapitulates the experimental results. Moreover, our model revealed that increasing the magnitude of variance resulted in increasing rates of desynchronization (**Supplementary Figure 2**). In order to further evaluate our model’s prediction, we sought to experimentally introduce an additional source of noise that might impact the overall variance of cell cycle duration.

### Impact of LPS on cell cycle duration variability

Lipopolysaccharide (LPS) is a major component of the outer membrane of Gram-negative bacteria that can bind to TLR4 receptors initiating a signaling cascade that ultimately results in NFkappaB translocation from the cytoplasm to the nucleus, where as a transcription factor, it initiates the upregulation of inflammation regulatory genes^16–18^. Additionally, NFkappaB activation can be induced by cytokines such as TNFalpha^19^, which has been reported with contrasting roles, whereby NFkappaB induction is associated with both the activation of pro-survival genes as well pro-apoptotic genes^20^. In addition to regulating inflammation signaling pathways, NFkappaB regulates major cell cycle regulatory factors^21–24^. Interestingly, components of NF-κB, such as RelA, have shown to interact with key cell cycle regulators, such as E2F transcription factors that are crucial in controlling progression through the G_1_/S boundary^22^.

We therefore hypothesized that the contrasting nature of LPS stimulation in HeLa cells would result in a greater variance in overall cell cycle duration. Accordingly, if LPS is a viable approach for introducing cellular noise we would expect the desynchronization rate to increase compared to untreated synchronized cells (**Supplementary Figure 3**). Thus, in order to determine if LPS simulation had any effect on cell cycle duration, we conducted a time-lapse experiment to track individual cells cell cycle duration. In order to have a better indication of relative position of each cell in relation to the cell cycle, we integrated a fluorescence tracker using lentiviral transduction that express the histone protein H2B fused to a fluorescent protein (H2B-FT)^25^. Upon expression, the H2B protein is incorporated into nucleosomes, which binds DNA, and therefore could more easily distinguish cells undergoing mitosis. Next, we treated asynchronously-growing HeLa cells with 1.0 μg/mL of LPS derived from *E. coli*. O111:B4, and monitored the duration of the cell cycle for individual cells with timelapse microscopy for 72 hours every 20 minutes (**Supplementary Figure 4, Supplementary Videos**). We found that the overall variance was higher in treated cells versus untreated cells with an accompanying increase in the mean duration 23.7±4.73 and 21.7±3.42 hours, respectively (**Figure 4a**). Additionally, we also found that the addition of LPS appeared to enhance cell motility, as LPS treated cells showed an increase in overall cell displacement (**Supplementary Figure 5**), which supports the notion that LPS can induce epithelial-mesenchymal transition via TLR4 signaling^26–28^.

**Figure 4:**
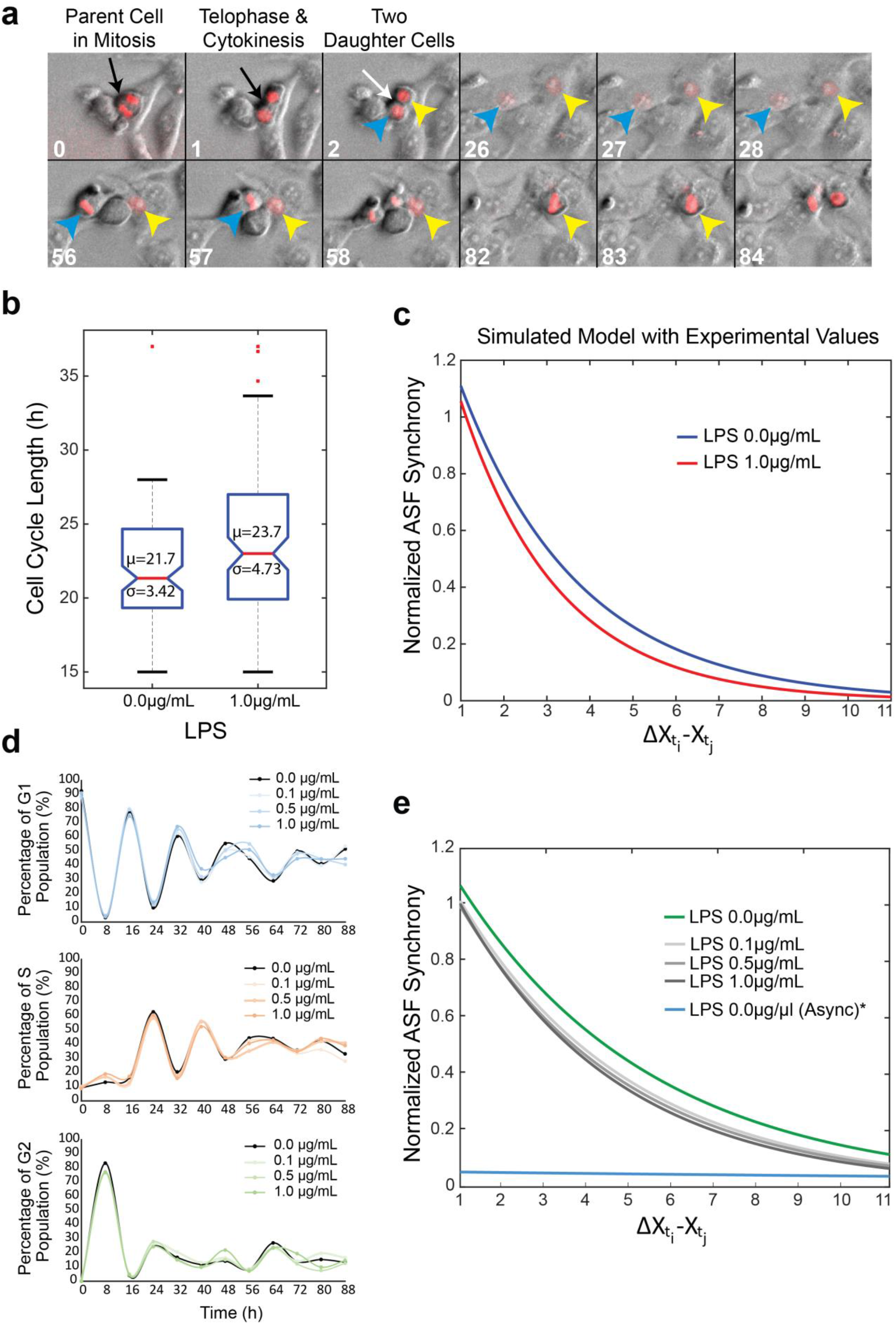
Noise variation of cell periodicity. **a)** Representative images of time lapse experiments. Once the septum (white arrow) is visible following cytokinesis, the cell cycle duration recording begins for both daughter cells (yellow and blue arrow). Both cells being cell cycle at Frame 2, and both daughter cells can be seen progressing through interphase in Frames 26-28. By the end of Frame 57, the first daughter cell completes the cell cycle and recording ends. The second daughter cell (yellow arrow) had a substantially longer cell cycle duration, which concluded at the end of Frame 84, thus demonstrating the inherent variability of cell cycle duration between identical cells within the population. **b)** Asynchronous cells were wither treated with 1 μg/mL of LPS or left untreated and cell cycle duration was recorded as described in Figure 4a. 100 cells were tracked for each condition and the population mean and standard deviation of cell cycle duration is shown. **c)** Values obtained from time lapse microscopy for cell cycle mean and standard deviation were used in our model to predict the impact on cell cycle desynchronization. The model revealed the LPS administration should result in an increased rate of cell cycle desynchronization **d)** Cell cycle phase distribution of various LPS concentrations following cell cycle synchronization for 88 hours post release. **e)** Normalized ASF scores for LPS-treated desynchronizing cells. The asynchronous population was not normalized in order to capture the overall linear trend.

We next sought to test if the predictability of our simulation model with the values obtained from the time-lapse microscopy would result in an increased desynchronization rate under LPS stimulation. In order to compare multiple synchronous cell samples, we normalized each sample to its initial ASF value (**X**_*t*_0__ – **X**_*t*_8__, **Supplementary Figure 6**). Upon inputting our new values obtained from the time-lapse microscopy, our model indeed predicted an increase in desynchronization when treated with LPS compared to the untreated sample (**Figure 4b**).

Given that we were able to increase the variance of cell cycle duration with LPS, and that our simulation model predicted an increase in desynchronization due to increased cell cycle duration variance, we next tested if we could experimentally obtain higher rates of desynchronization using the previous approach of PI-staining time-separated synchronized cells. Therefore, we again synchronized HeLa cells via the double-thymidine block method, and immediately following release of the arrested cells, we treated with varying concentrations of LPS (0.0, 0.1, 0.5, 1.0 μg/mL) and collected timepoints every 8 hours for 88 hours. We then analyzed the PI-stained cell populations via FlowJo cell cycle classifier that uses the Dean-Jett-Fox algorithm, and while it was not able to detect substantial changes in the overall phase distributions at the early timepoints (0-32 hours), the effects of LPS were more evident at the later timepoints (40-88 hours) (**Figure 4c**). Interestingly, our ASF analysis methodology (**Supplementary Figure 7-8**) was able to differentiate the impact on cell cycle desynchronization at all timepoints tested for all LPS concentrations, with each increase in LPS having a greater impact in cell cycle desynchronization (**Figure 4d**).

## Discussion

The cell cycle and subsequent daughter cell division is a central facet of cell biology from development and cellular differentiation to disease initiation and progression. While the process is tightly regulated and robust in a single cell, across a population we observe significant variability in period. Each cell within a given population contains measurable variations in their cellular content and housekeeping genes (e.g., differences in their RNA polymerases, ribosomes). These variations impact the expression of genes in what is known as extrinsic noise^29–33^. Furthermore, the cellular machinery responsible for progressing a cell through its cycle is intrinsically noisy as well. These intra and inter-cellular differences cause an initially synchronously in-phase population of cells to diverge as each progresses independently through their life cycle at varying rates^34^.

Moreover, cell synchronicity is an essential aspect of mammalian biological homeostasis. The circadian rhythm is a molecular orchestrated process present in various tissues that synchronizes biological outputs to the 24-hour day-night cycle^35^. It is composed of multiple master transcription factor regulators that are involved in robust feedback networks that enable these oscillatory functions is cells^36^. Importantly, cancer cells can become compromised with respect to these molecular actuators the enable cellular synchronization, and there is evidence that this disruption can enhance tumorigenesis^37^. For example, DNA repair mechanisms are tied to the circadian regulation^38^ and if these processes are disrupted it may lead to accumulating mutations and overall genome instability^39^. Thus, understanding and modeling cell desynchronization at a single-cell level is a decisive approach to quantifying cellular changes that may impact oncogenesis^40^.

Here, using a combination of simulations and experiments we show that the variability in cell cycle period directly impacts the rate of desynchronization in a population of cells. The next line of investigation will include studying the factors that contribute to this variability at a single cell level, and the distribution between intrinsic and extrinsic sources of noise. Finally, a key direction of research is not just on how such a tightly regulated process can rely on stochastic variations, but also how diseased states such as cancer cells, utilize noise and if noise itself can become a contributor to disease progression. More specifically an intriguing hypothesis is that cancer cells obtain benefit by having higher noise in cell cycle periodicity, which yields ultra-slow and fast diving cells. Moreover, this hypothesis opens the path for potential means to exploit variability in cell cycle period for therapeutic purposes.

## Methods and Procedures

### Cell culturing and synchronization

HeLa cells were grown in Gibco DMEM supplemented with 10%FBS, 1X PenStrep, 2mM glutamine, and 1X Gibco NEAA and grown at 37°C with 5% CO_2_. 50,000 cells were seeded per well in 6 well plates. 24 hours post-seeding cells were treated with 2mM of thymidine for 19 hours after which the cells were washed with 1X PBS and given fresh complete media to release from the first thymidine block. The cells then incubated for 9 hours before receiving a second dose of 2mM of thymidine for 15 hours. Cells were washed with 1X PBS to remove thymidine before given fresh media to continue to grow unimpeded. Cells harvested at t=0 were collected immediately following the second PBS wash. Additional wells were harvested every 8 hours for 88 hours. Asynchronous cells were harvested at same time as synchronized cells for each time point. Cells were harvested by washing with PBS, detached from the well with trypsin-EDTA (0.25%) for 3 min at 37°C then quenched with fresh complete media. Harvested cells were pelleted at 1000rpms for 5 min at room temperature. The supernatant was removed and the cell pellet was resuspended in 1X PBS, then pelleted again at 1000rpms for 5 min at room temperature. The supernatant was removed and the cell pellet was resuspended in 1 mL of 70% ethanol and stored at 4°C for a minimum of 24 hours to fix the cells.

LPS derived from *E. coli* 0E111 was reconstituted in PBS without Mg or Ca at a concentration of 1mg/mL. LPS solution was added directly to the cell culture media after replacing with fresh media initiating the release from the double thymidine arrested state.

### Propidium Iodide Staining

After fixation, cells were pelleted by centrifuged at 1000 rpms for 5 minutes at room temperature. The fixing solution was aspirated off the cell pellet, and resuspended in 1X PBS. Cells were counted for each sample, and then normalized to the lowest cell count for uniform propidium iodide (PI) staining across samples. The PI staining procedure was done according to manufacturer’s directions (Propidium Iodide Flow Cytometry Kit, cat# ab139418).

### Cell Cycle Phase Analysis

Stained cells were subjected flow cytometry using a BD LSRFortessa^™^ flow cytometer. PI fluorescence was excited with a 561nm laser and emission was detected using a 610/20 nm band-pass filter. Assignment of cell cycle phases were performed using the univariate modeling via the Dean-Jett-Fox algorithm with FlowJo 10.7.1.

### Lentiviral HeLa Transduction for H2B-FT expression

The fluorescent tracker sequence was obtained from addgene (#157671) and cloned using primers P1: gaagagttcttgcagctcggtgac and P2: cagtagggtaccccggaattagatcgatctctcgacatcc. The amplicon was digested with restriction enzymes BsiWI and KpnI and inserted into the LentiCRISPRv2 (addgene #52961) backbone. The resulting plasmid was transfected into HEK293T cells along with pMD-VSVG and psPAX2 plasmids to generate viral particles that are released into the media. The media was aspirated two days post-transfection, and replenished with 5 mL of fresh media every day for three days. The 15 mL of harvested viral-containing media was passed through a 0.45 μm filter and dispensed into 1 mL aliquots. 250 μl was used to transduce HeLa cells, and 0.5μg/mL of Puromycin was used to select for integrated clones for 7 days.

### Time-lapse Microscopy

Images were collected every 20 min for 72 hours using Hamamatsu camera attached to the Olympus IX81 microscope at 10x magnification. Cells were maintained at 37°C and 5% CO_2_. The exposure time was 250 ms for Brightfield and 100ms for TexasRed using Chroma filter ET560/40x (excitation) and ET630/75m (emission).

### Modeling and Simulations

All models and simulations were developed and tested in Mathematica.

## Acknowledgements

We thank Khai Nguyen and Bleris lab members for support and discussions. This work was funded by US National Science Foundation (NSF) grants (1351354, 1361355, 2029121), a Cecil H. and Ida Green Endowment, and the University of Texas at Dallas.

## Author contributions

C.M.N. performed the experiments. C.M.N., T.Q., and L.B. analyzed the data and developed models. C.M.N., T.Q., and L.B. wrote the paper. L.B. supervised the project.

No competing financial interests

## Additional Information

Supplementary information is available in the online version of the paper. Correspondence and requests for materials should be addressed to L.B.

## References

1. Smith, J. A. & Martin, L. Do Cells Cycle? (cell kinetics/control of growth/DNA replication/cell culture). DETERMINATE Downloaded at LIBRARY SERIALS vol. 70 (1973).

2. Prescott, D. M. Regulation of Cell Reproduction. Cancer Res. 28, (1968).

3. Temin, H. M. Stimulation by serum of multiplication of stationary chicken cells. J. Cell. Physiol. 78, 161–170 (1971).

4. Pardee, A. B. A Restriction Point for Control of Normal Animal Cell Proliferation (growth control/cell survival/cancer). vol. 71 (1974).

5. Zetterberg, A. & Larsson, O. Kinetic analysis of regulatory events in G1 leading to proliferation or quiescence of Swiss 3T3 cells (GI arrest/GO state/epidermal growth factor/platelet-derived growth factor/insulin). Cell Biology vol. 82 (1985).

6. Schorl, C. & Sedivy, J. M. Analysis of Cell Cycle Phases and Progression in Cultured Mammalian Cells. Methods 41, 143 (2007).

7. PK, D., A, H. & SF, D. Biological methods for cell-cycle synchronization of mammalian cells. Biotechniques 30, 1322–1331 (2001).

8. Bjursell, G. & Reichard, P. Effects of thymidine on deoxyribonucleoside triphosphate pools and deoxyribonucleic acid synthesis in Chinese hamster ovary cells. J. Biol. Chem. 248, 3904–3909 (1973).

9. Chen, G. & Deng, X. Cell Synchronization by Double Thymidine Block. BIO-PROTOCOL 8, (2018).

10. Chiorino, G., Metz, J. A. J., Tomasoni, D. & Ubezio, P. Desynchronization rate in cell populations: Mathematical modeling and experimental data. J. Theor. Biol. 208, 185–199 (2001).

11. Nguyen, A., Yoshida, M., Goodarzi, H. & Tavazoie, S. F. Highly variable cancer subpopulations that exhibit enhanced transcriptome variability and metastatic fitness. Nat. Commun. 7, (2016).

12. O’Duibhir, E. et al. Cell cycle population effects in perturbation studies. Mol. Syst. Biol. 10, 732 (2014).

13. Watson, J. V., Chambers, S. H. & Smith, P. J. A pragmatic approach to the analysis of DNA histograms with a definable G1 peak. Cytometry 8, 1–8 (1987).

14. Fox, M. H. A model for the computer analysis of synchronous DNA distributions obtained by flow cytometry. Cytometry 1, 71–77 (1980).

15. Dean, P. N. & Jett, J. H. Mathematical analysis of dna distributions derived from flow microfluorometry. J. Cell Biol. 60, 528–527 (1974).

16. Wang, Y. et al. Expression and Functional Analysis of Toll-like Receptor 4 in Human Cervical Carcinoma. J. Membr. Biol. 2014 2477 247, 591–599 (2014).

17. Savinova, O. V., Hoffmann, A. & Ghosh, G. The Nfkb1 and Nfkb2 Proteins p105 and p100 Function as the Core of High-Molecular-Weight Heterogeneous Complexes. Mol. Cell 34, 591 (2009).

18. N, J. et al. Toll-like receptor 4 promotes proliferation and apoptosis resistance in human papillomavirus-related cervical cancer cells through the Toll-like receptor 4/nuclear factor-κB pathway. Tumour Biol. 39, (2017).

19. Hayden, M. S. & Ghosh, S. Regulation of NF-κB by TNF Family Cytokines. Semin. Immunol. 26, 253 (2014).

20. Lee, R. E. C., Qasaimeh, M. A., Xia, X., Juncker, D. & Gaudet, S. NF-κB signalling and cell fate decisions in response to a short pulse of tumour necrosis factor. Sci. Reports 2016 61 6, 1–12 (2016).

21. Bash, J., Zong, W. X. & Gélinas, C. c-Rel arrests the proliferation of HeLa cells and affects critical regulators of the G1/S-phase transition. Mol. Cell. Biol. 17, 6526 (1997).

22. Ankers, J. M. et al. Dynamic NF-κb and E2F interactions control the priority and timing of inflammatory signalling and cell proliferation. Elife 5, (2016).

23. Ledoux, A. C. & Perkins, N. D. NF-κB and the cell cycle. Biochem. Soc. Trans. 42, 76–81 (2014).

24. Kenter, A. L. & Watson, J. V. Cell cycle kinetics model of LPS-stimulated spleen cells correlates switch region rearrangements with S phase. J. Immunol. Methods 97, 111–117 (1987).

25. Eastman, A. E. et al. Resolving Cell Cycle Speed in One Snapshot with a Live-Cell Fluorescent Reporter. Cell Rep. 31, 107804 (2020).

26. Jing, Y.-Y. et al. Toll-like receptor 4 signaling promotes epithelial-mesenchymal transition in human hepatocellular carcinoma induced by lipopolysaccharide. BMC Med. 2012 101 10, 1–12 (2012).

27. Li, H., Li, Y., Liu, D. & Liu, J. LPS promotes epithelial–mesenchymal transition and activation of TLR4/JNK signaling. Tumor Biol. 2014 3510 35, 10429–10435 (2014).

28. Cho, I.-H. et al. Suppression of LPS-induced epithelial-mesenchymal transition by aqueous extracts of Prunella vulgaris through inhibition of the NF-κB/Snail signaling pathway and regulation of EMT-related protein expression. Oncol. Rep. 34, 2445–2450 (2015).

29. Quarton, T. et al. Uncoupling gene expression noise along the central dogma using genome engineered human cell lines. Nucleic Acids Res. 48, 9406–9413 (2020).

30. Elowitz, M. B., Levine, A. J., Siggia, E. D. & Swain, P. S. Stochastic gene expression in a single cell. Science (80–.). 297, 1183–1186 (2002).

31. Kang, T. et al. Robust Filtering and Noise Suppression in Intragenic miRNA-Mediated Host Regulation. iScience 23, 101595 (2020).

32. Raser, J. M. & O’Shea, E. K. Control of stochasticity in eukaryotic gene expression. Science (80–.). 304, 1811–1814 (2004).

33. Swain, P. S., Elowitz, M. B. & Siggia, E. D. Intrinsic and extrinsic contributions to stochasticity in gene expression. Proc. Natl. Acad. Sci. 99, 12795–12800 (2002).

34. Chao, H. X. et al. Evidence that the human cell cycle is a series of uncoupled, memoryless phases. Mol. Syst. Biol. 15, e8604 (2019).

35. Patke, A., Young, M. W. & Axelrod, S. Molecular mechanisms and physiological importance of circadian rhythms. Nat. Rev. Mol. Cell Biol. 2019 212 21, 67–84 (2019).

36. Partch, C. L., Green, C. B. & Takahashi, J. S. Molecular Architecture of the Mammalian Circadian Clock. Trends Cell Biol. 24, 90 (2014).

37. Sulli, G., Lam, M. T. Y. & Panda, S. Interplay between circadian clock and cancer: new frontiers for cancer treatment. Trends in cancer 5, 475 (2019).

38. Sancar, A. et al. Circadian Clock Control of the Cellular Response to DNA Damage. FEBS Lett. 584, 2618 (2010).

39. Gery, S. et al. The Circadian Gene Per1 Plays an Important Role in Cell Growth and DNA Damage Control in Human Cancer Cells. Mol. Cell 22, 375–382 (2006).

40. Barberis, M. & Verbruggen, P. Quantitative Systems Biology to decipher design principles of a dynamic cell cycle network: the “Maximum Allowable mammalian Trade–Off–Weight” (MAmTOW). npj Syst. Biol. Appl. 2017 31 3, 1–14 (2017).

